# Tessellation of artificial touch via microstimulation of human somatosensory cortex

**DOI:** 10.1101/2023.06.23.545425

**Authors:** Charles M. Greenspon, Natalya D. Shelchkova, Giacomo Valle, Taylor G. Hobbs, Ev I. Berger-Wolf, Brianna C. Hutchison, Efe Dogruoz, Ceci Verbarschott, Thierri Callier, Anton R. Sobinov, Elizaveta V. Okorokova, Patrick M. Jordan, Dillan Prasad, Qinpu He, Fang Liu, Robert F. Kirsch, Jonathan P. Miller, Ray C. Lee, David Satzer, Jorge Gonzalez-Martinez, Peter C. Warnke, Lee E. Miller, Michael L. Boninger, Abidemi B. Ajiboye, Emily L. Graczyk, John E. Downey, Jennifer L. Collinger, Nicholas G. Hatsopoulos, Robert A. Gaunt, Sliman J. Bensmaia

## Abstract

When we interact with objects, we rely on signals from the hand that convey information about the object and our interaction with it. A basic feature of these interactions, the locations of contacts between the hand and object, is often only available via the sense of touch. Information about locations of contact between a brain-controlled bionic hand and an object can be signaled via intracortical microstimulation (ICMS) of somatosensory cortex (S1), which evokes touch sensations that are localized to a specific patch of skin. To provide intuitive location information, tactile sensors on the robotic hand drive ICMS through electrodes that evoke sensations at skin locations matching sensor locations. This approach requires that ICMS-evoked sensations be focal, stable, and distributed over the hand. To systematically investigate the localization of ICMS-evoked sensations, we analyzed the projected fields (PFs) of ICMS-evoked sensations – their location and spatial extent – from reports obtained over multiple years from three participants implanted with microelectrode arrays in S1. First, we found that PFs vary widely in their size across electrodes, are highly stable within electrode, are distributed over large swaths of each participant’s hand, and increase in size as the amplitude or frequency of ICMS increases. Second, while PF locations match the locations of the receptive fields (RFs) of the neurons near the stimulating electrode, PFs tend to be subsumed by the corresponding RFs. Third, multi-channel stimulation gives rise to a PF that reflects the conjunction of the PFs of the component channels. By stimulating through electrodes with largely overlapping PFs, then, we can evoke a sensation that is experienced primarily at the intersection of the component PFs. To assess the functional consequence of this phenomenon, we implemented multi-channel ICMS-based feedback in a bionic hand and demonstrated that the resulting sensations are more localizable than are those evoked via single-channel ICMS.

## Introduction

When we interact with objects, signals from the hand convey information about the objects and about our interactions with them (Johansson and Flanagan, 2009). A basic feature of object interactions, the location on the hand of object contacts, is often only available via the sense of touch because some contacts are visually occluded and visual feedback, even when available, is a poor substitute for touch to guide object interactions (Cabibihan et al., 2021). Efforts are underway to restore tactile feedback via bionic hands by electrically activating touch neurons along the neuraxis (Bensmaia et al., 2020). Indeed, electrical activation of tactile nerve fibers evokes sensations experienced at a specific location on the skin, known as the projected field (PF) (Ochoa and Torebjörk, 1983; Ortiz-Catalan et al., 2020b; Overstreet et al., 2019; Petrini et al., 2019; Tan et al., 2014; Wendelken et al., 2017). Similarly, intracortical microstimulation (ICMS) of somatosensory cortex (S1) typically evokes a sensation restricted to a specific patch of skin (Fifer et al., 2020; Flesher et al., 2016; Salas et al., 2018; Tabot et al., 2013). Force sensors on the bionic hand can thus be used to drive stimulation through electrodes in the nerve or in the brain that evoke sensations on the corresponding location on the phantom or deafferented hand, thereby intuitively conveying information about contact location.

The representation of the body in S1 is systematically organized, such that nearby body parts are encoded by the activity recorded from neighboring neuronal populations, with some discontinuities reflecting those in the body (Delhaye et al., 2018; Penfield and Boldrey, 1937). Stimulating through nearby electrodes has been shown to evoke sensations in nearby hand regions following the expected somatotopic organization (Fifer et al., 2020; Flesher et al., 2016). Furthermore, the organization of S1 is consistent across individuals – with the hand representation featuring a systematic progression from the thumb to the little finger as one progresses in the medial posterior direction along the central sulcus. This organization facilitates electrode array placement because locating the respective S1 representations of a subset of digits informs the localization of the others. Pre-operative functional magnetic resonance imaging (fMRI), magnetoencephalography (MEG), or electroencephalography can be used to identify the digit representations and thus guide the placement of arrays of stimulating electrodes (Flesher et al., 2016; Foldes et al., 2021; Herring et al., 2023). Critically, studies involving limited numbers of patients and electrodes have shown that PFs are relatively stable over time (Flesher et al., 2016; Ortiz-Catalan et al., 2020a), suggesting that the somatotopic map is stable, even after deafferentation caused by spinal cord injury or amputation (Kikkert et al., 2021, 2016; Makin and Bensmaia, 2017).

Though electrical activation of S1 neurons has been shown to evoke focal sensations, these observations have been largely qualitative. The spatial extent, distribution, and stability of the evoked PFs have not been systematically evaluated (Bjånes et al., 2022; Flesher et al., 2016; Salas et al., 2018). To fill this gap, we quantitatively characterized the PFs of ICMS-evoked sensations in three human participants with cervical spinal cord injury and gauged the degree to which they remained stable over a period of years. In an additional series of experiments with participants endowed with residual sensation, we also assessed the degree to which the PF of an electrode coincided with its receptive field (RF), defined as the patch of skin that, when touched, activates neurons around the electrode tip. First, we found that PFs are highly stable over time and cover large swaths of the hand (across two arrays in S1). Second, the locations of the PFs follow the expected somatotopic organization of the sensory homunculus. Third, increasing the amplitude or frequency of ICMS increases the size of the PF. Fourth, at the current levels tested, the PF of an electrode tends to be smaller than but largely subsumed by its RF. Fifth, ICMS delivered through multiple electrodes yields a PF reflecting a conjunction of the PFs of the individual electrodes; any overlapping regions of the PFs are thus more salient. Accordingly, stimulation through multiple electrodes with overlapping PFs enables improved localization of touches on a bionic hand compared to single-electrode stimulation. We discuss the implications of our findings for the development of sensitized brain-controlled bionic hands.

## Results

Three participants were implanted with arrays of electrodes in the hand representation of S1 (Brodmann’s area 1, **Figure 1, Supplementary Figure 1**), localized based on pre-operative fMRI or MEG. In participant C1, most of the sensations were experienced on the fingertips, in participant P2, on the palm, and in participant P3, on the medial phalanges and palm. To quantitatively characterize the PF of each electrode, we repeatedly delivered through it a 1-sec long ICMS pulse train (100 Hz, 60 μA), which was intense enough to reliably elicit a sensation for most electrodes (**Supplementary Figure 2**). After each stimulation bout, the participant drew the spatial extent of the PF on a digital representation of his hand, enabled by partial residual arm function. From these images, we computed the area of skin over which the PF extended and its center of mass (centroid) (**Figure 2A**). This task was repeated regularly over several years (**Figure 2B**), allowing us to construct an aggregate PF for each stimulating channel by weighting each pixel on the hand by the proportion of times it was included in the reported PF over the duration of the study (**Figure 2C**). This allowed us to estimate the probability of a sensation being evoked on different parts of the hand for each channel and revealed that most PFs comprised a core region within which sensations were reliably evoked, surrounded by a diffuse shell over which sensations were less consistent. With this in mind, we applied a reliability criterion (33%) on the aggregate PFs and removed pixels that did not meet this criterion (see **Supplementary Figure 3** for the justification of the threshold level). PFs for which no pixels met this threshold criterion were deemed too unreliable and excluded from further analysis, totaling around 25% for Participants C1 & P2 and ∼60% for Participant P3. While the excluded PFs were typically reported on the same digit or hand region (**Figure 2 B, C** - green), we reasoned that they were not sufficiently reliable to usefully convey location information.

**Figure 1.**
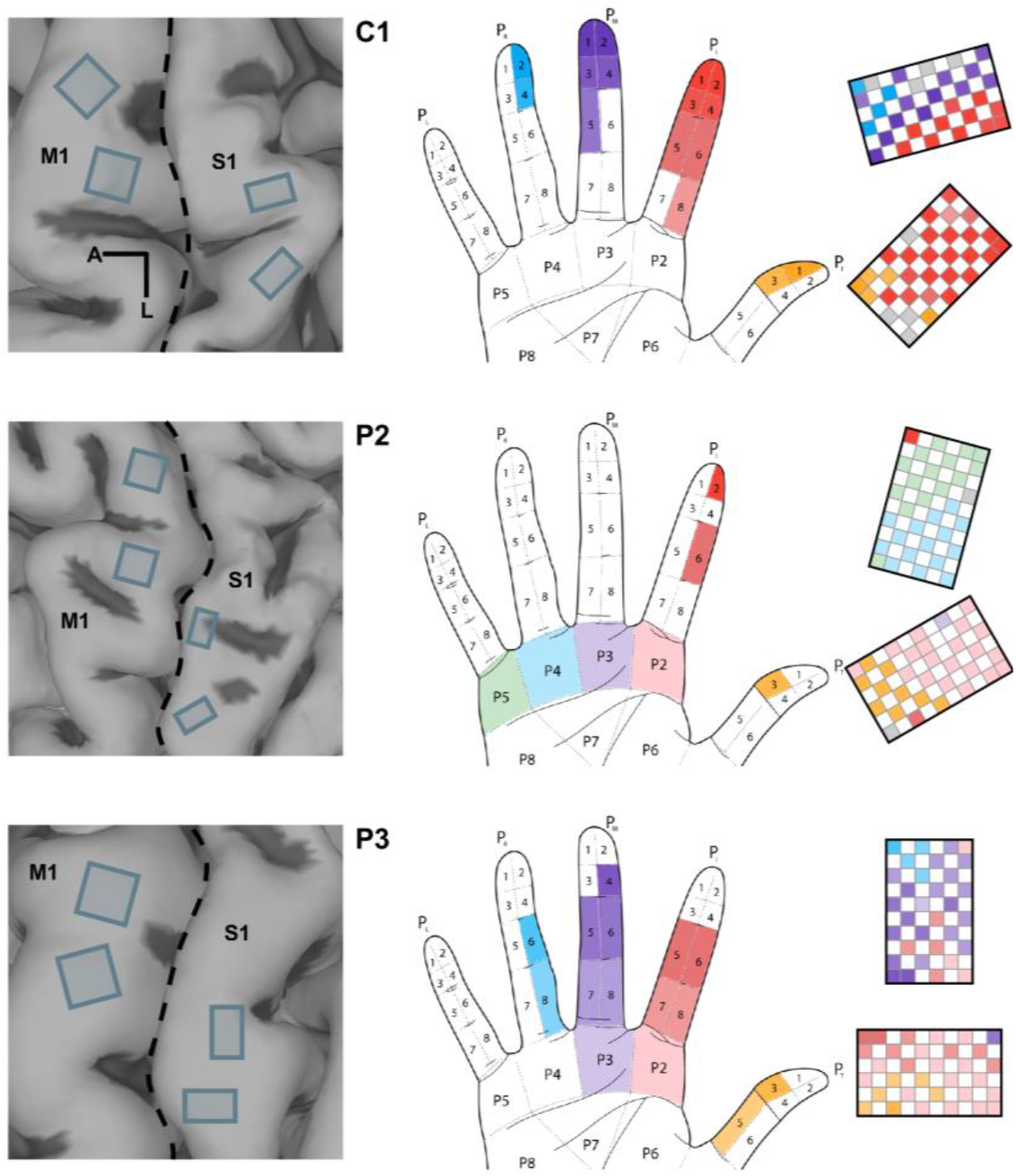
Array implant locations and sensation maps for all participants. *Left column*: Anatomical MRI with (subsequently implanted) arrays superimposed whose location is based on intra-operative photos. M1 and S1 denote primary motor cortex and somatosensory cortex (Brodmann’s area 1), respectively. The central sulcus is indicated by the dashed line. *Middle and right columns*: The hand region on which ICMS-sensations are experienced along with the electrodes that evoked those sensations. Each row shows data from one participant.

**Figure 2.**
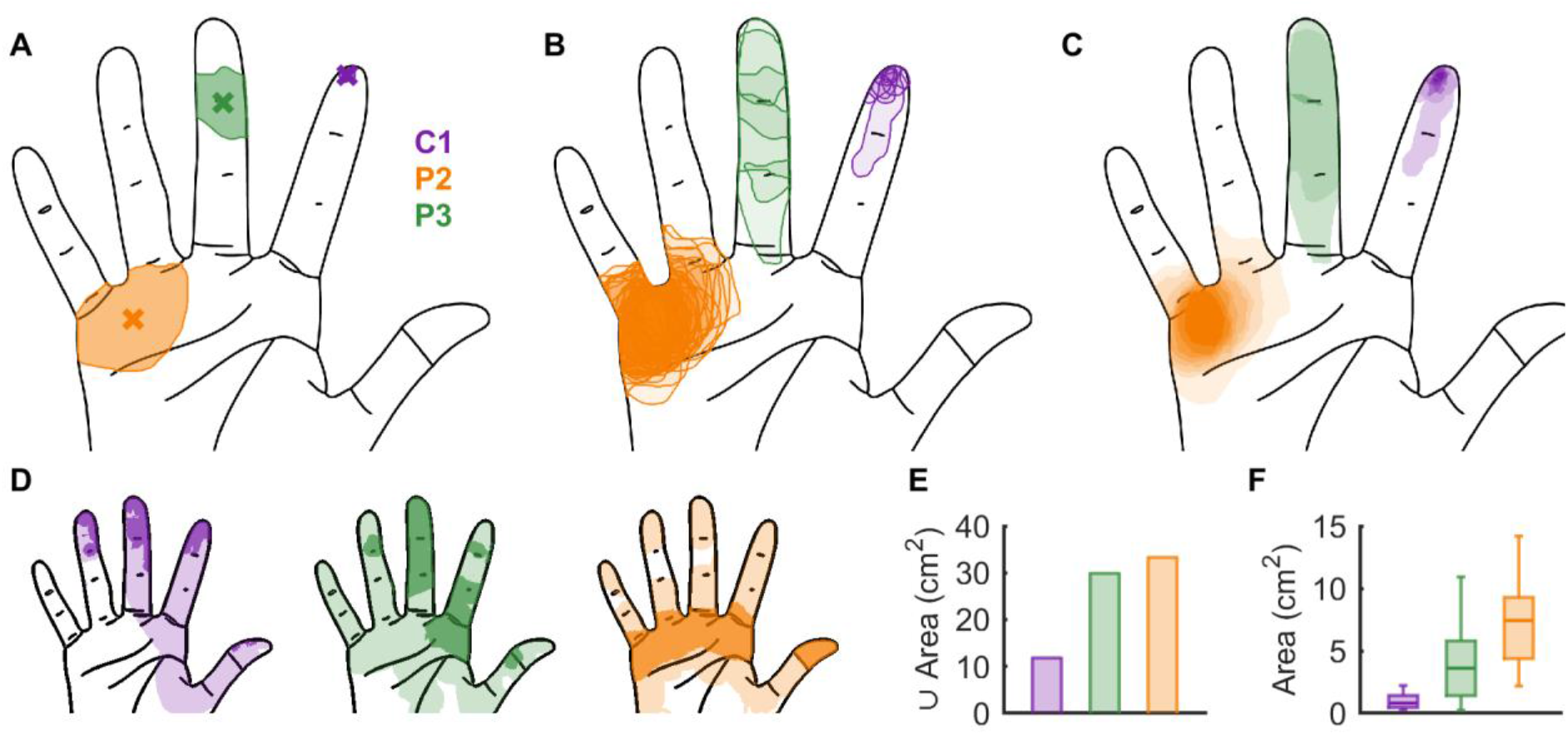
Projected field locations systematically vary in size and location across electrodes and participants. **A**| Example projected fields from one session from each participant. Crosses denote the respective centroids. **B|** The same projected fields as in panel A but across all sessions. **C|** The density function computed for the same electrodes across all sessions. **D|** Hand regions over which each participant reported a sensation across all electrodes. The lighter shade indicates pixels selected on <33% of sessions while the darker shade indicates pixels selected on >33% of sessions. **E|** The area of the hand over which a sensation was evoked (union of PFs across electrodes, after thresholding) for each participant. **F|** The distribution of individual PF sizes (after thresholding) for each participant.

Next, we examined how the PFs tiled the hands for each participant. While all participants reported a sensation over most of the hand on at least one occasion (**Figure 2D**, light shade), participant C1 predominantly experienced sensations on the tip of the digits, participant P2 on the pads and thumb, and participant P3 on both. The total area over which sensations were evoked (after thresholding) was 11.8, 33.3, and 29.9 cm^2^ for C1, P2, and P3, respectively (**Figure 2E**). The size of the PF varied widely (median = 2.5 cm^2^; 5^th^/95^th^ percentiles = 0.3 / 11.3 cm^2^) across electrodes and participants (**Figure 2F**), with C1 reporting the smallest PFs and P2 the largest. C1 had the most distal PFs and P2 had the most proximal ones, suggesting that PFs decrease in size as one progresses distally from the palm to the digit tips but confounded by the small sample of participants. Note that receptive fields (RFs) of neurons in S1 generally follow an analogous trend, where distal RFs are smaller than proximal ones (Sur et al., 1980). Similarly, the size of ICMS-evoked phosphenes via ICMS of primary visual cortex increases as one proceeds toward the visual periphery (Bosking et al., 2017).

### Projected fields are highly stable over time

While PF field location has been reported to be stable over time (Flesher et al., 2016), stability has not been systematically quantified. To fill this gap, we compared the location of the PFs at regular intervals across 2 to 7 years (for C1/P3 and P2, respectively; **Figure 2B,C**). First, we computed the degree to which PFs reported for an electrode on any given session matched the first ever reported PF on that electrode. We found that centroid distance between the initial PF and subsequent ones for a given electrode remained stable, with no significant change over the lifetime of the arrays for two of the three participants (C1 and P3, r = -0.03 and 0.12, p = 0.99 and 0.34, respectively), and a slight but significant increase in the third participant, who had been implanted the longest (P2, r = 0.23, p < 0.01, **Supplementary Figure 4A**). In other words, we observed little to no progressive shift away from the first reported PF with subsequent reports. At the single electrode level, the slopes of the PF shifts over time (in mm per day) – some negative denoting trends toward the first reported PF after an initial increase – were always very shallow (median = 0.00 mm/day; Q_1_, Q_3_ = 0.00, 0.01 mm/day respectively) and their mean was only significantly different from zero for one participant (P2, t(58) = 3.35, p < 0.01, **Supplementary Figure 4B**). Second, we computed the distance between each single-day centroid and its corresponding aggregate centroid (**Figure 3A**) to determine how representative the aggregate PF was of any individual PF for a given electrode. Across all recording sessions, we found that the mean distance ranged from 3.5 to 8.7 mm (depending on the participant), but larger PFs tended to yield more variable centroids (**Figure 3B**, r = 0.33, p < 0.01), as might be expected. Furthermore, centroid distance was stable over time for one participant (P3, r = -0.1, p = 0.77) and *decreased* over time for the other two (C1 and P2, r = -0.17 for both, p < 0.01 for both), suggesting that the latter converged onto a more stable reporting of their PFs (over the years that these were measured).

**Figure 3.**
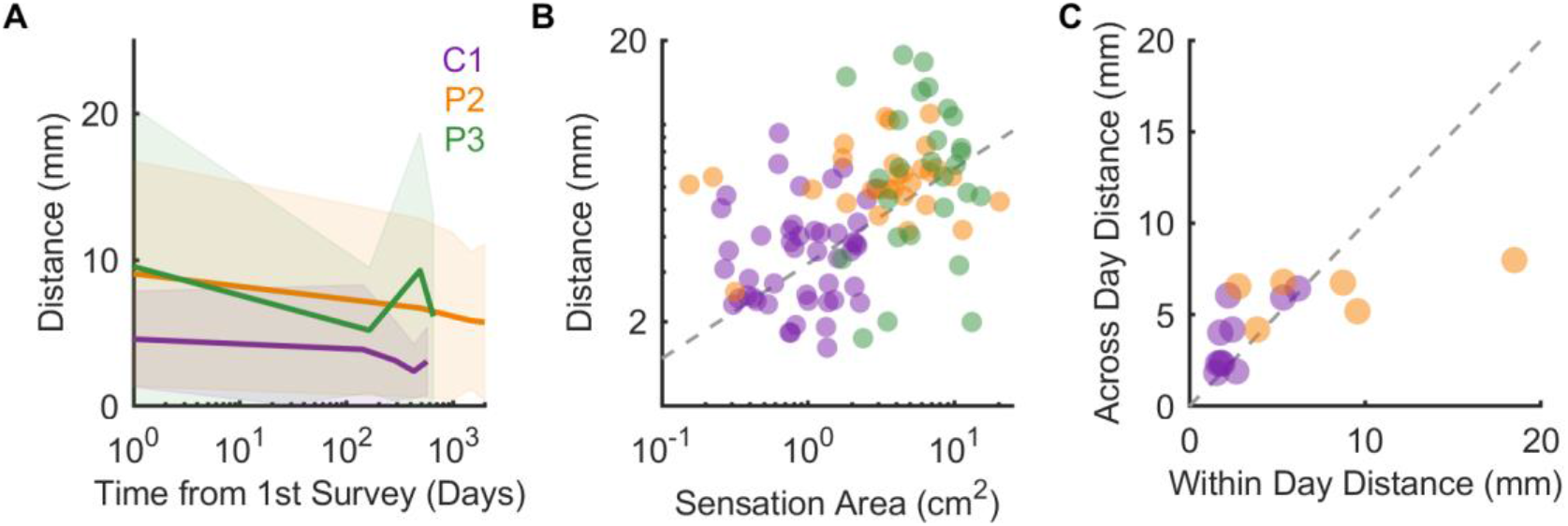
Projected field locations are stable over time. **A**| Distance between the centroid of the single-day PF and the aggregate centroid for each electrode, averaged across electrodes. The line denotes the mean and the shaded area the standard deviation. **B|** Mean distance between single day PF centroid and aggregate PF centroid for each electrode. Mean distance increases with the size of the PF. Dashed line denotes best fit. **C|** Mean centroid distance when reports were collected within a single day compared to that computed across years for a subset of electrodes. Dashed line denotes unity.

Next, we assessed whether the fluctuations in reported PF reflected true changes in position over time or simply variability in the participants’ reports. To this end, ICMS pulse trains were delivered through a subset of electrodes (n = 8 and 7 for C1 and P2, respectively), interleaved randomly, and the participant reported the PF of each. This procedure was then repeated multiple times throughout the day (6 and 5 times for participants C1 and P2, respectively), each time in a different order to reduce biases. We then assessed the degree to which the centroids of the reported PFs coincided. First, PFs were consistent between repeated reports for the two participants tested (centroid distance: median = 3.5 mm, Q_1_-Q_3_, 1.6 – 4.8 mm). Second, the within-vs. across-day centroid distances for each electrode were highly correlated (r = 0.68, p < 0.01) and statistically equivalent (paired t-test, t(15) = 0.1, p = 0.93, **Figure 3C**). In other words, the change in PF reported within a session was equivalent to the change in PF reported over years of testing. We conclude that the bulk of the variability in PFs over time reflects variability in the reports rather than variability in the PFs themselves (assuming that PFs are stable within a day). The PF is thus highly stable over time and well described by the aggregate PF.

### Projected fields progress systematically with location along the cortical surface

According to the canonical homunculus, the representations of digits and hand segments in S1 proceed systematically along the mediolateral axis, approximately parallel to the central sulcus, with the little finger represented medially and the thumb represented laterally (Penfield and Boldrey, 1937). Examination of the PF maps (**Supplementary Figure 1**) suggests that, within each array, the dominant digit or hand segment – where the PFs are predominantly located – also changes systematically along one axis, consistent with the geometry of the homunculus. To test this quantitatively, we projected the location of each electrode on each array onto a single axis and assessed the degree to which the identity of the digit could be inferred from the electrode’s location along that axis (**Figure 4A**). By computing classification accuracy across a range of axes (spanning 180 degrees), we identified the axis along which the digit/hand segment gradients were most pronounced. We found that, for each array, a single axis could account for digit or hand segment dominance as well as if both coordinates were included (**Supplementary Figure 5A**) and that this axis was approximately parallel to the local curvature of S1 for 5 out of the 6 arrays (**Figure 4B, Supplementary Figure 5**), consistent with the canonical homunculus. In the array that did not show this pattern (participant P3’s lateral array), the axis was nearly perpendicular to the S1 curvature, but the PFs were largely confined to a single digit (the index finger), yielding comparable classification at all angles. Next, we examined whether the distance between PFs increased as the distance between electrodes increased, restricting the analysis to pairs of electrodes with PFs on the same digit. We found that, as expected, centroid distance increased systematically with cortical distance (**Figure 4C**). A 2-mm shift in cortex corresponded to a ∼10-mm shift in the PF on the skin, on average. The systematic relationship between cortical location and PF location is thus consistent with the canonical homunculus, in which S1 is characterized by strips that encode individual digits or palmar segments.

**Figure 4.**
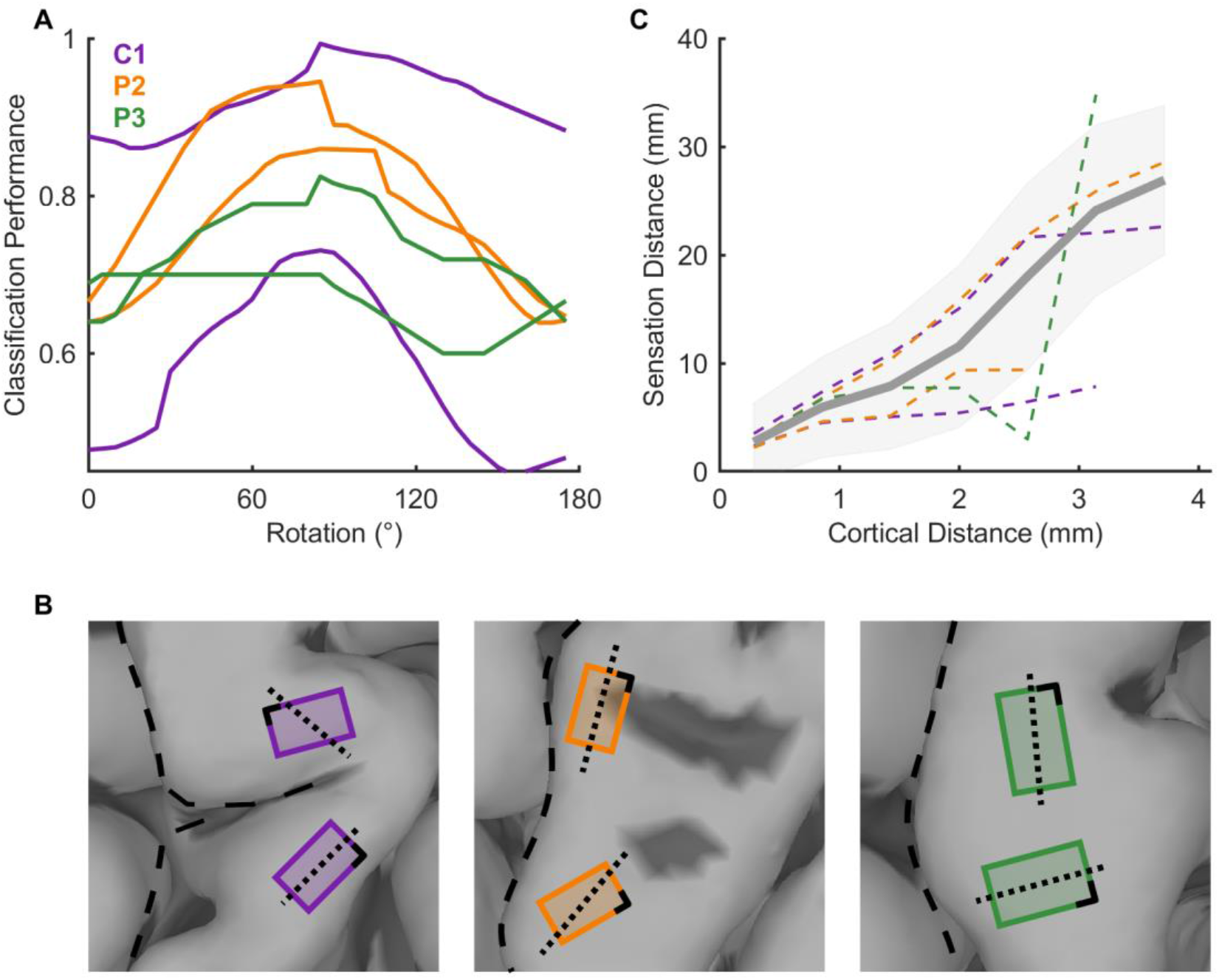
PFs progress systematically across electrodes. **A**| Classification performance vs. the angle of the projection axis, expressed relative to the local curvature of S1. The ability to infer digit or palmar segment identity based on position along a single axis depends on the angle of that axis. **B|** Optimal digit/palmar segment discrimination axis (perpendicular to the projection axis), superimposed on each S1 array (dotted line). The dashed line denotes the local curvature of S1, which, for C1, deviates from the curvature of the central sulcus. **C|** Within digit, the distance between two PF centroids is significantly correlated with the distance between the electrodes (n = 5892 pairs, r = 0.69, p < 0.01) and this relationship is observed for each array individually (r > 0.4, p < 0.01).

### The projected field is determined by the receptive field of the activated neurons

To further examine the relationship between homunculus location and PF location, we leveraged the residual sensation in two of our participants (C1 and P3, **Supplementary Figure 6**) and compared the RF and PF of each electrode. Indeed, C1 has nearly normal sensation over the fingertips and some residual sensation over the rest of the volar surface of his hand. P3 has largely spared sensation on the thumb, some spared sensation on the index, and some residual sensation on D3 and D4, but his little finger is insensate. To map RFs, we applied gentle touches to the skin and monitored the multi-unit activity through speakers. As for the PFs, RFs were measured over multiple days with the experimenter blinded to electrode identity. Repeated mapping sessions (in participant C1) yielded very similar RFs (see **Figure 5A left**). We then compared the RF and PF of each electrode (**Figure 5A**). First, we found that, across electrodes and participants, the RF was 6.5 times as large as the PF (**Figure 5B**, ratio: Q_2_ = 6.5, Q_1_-Q_3_ = 3.8 – 14.3) and PF size tended to scale with RF size, though this relationship was only significant for one of the two participants (r = 0.15 and 0.76, p = 0.35 and 0.011 for C1 and P3, respectively). Secondly, PFs were typically subsumed by the RF: On average, 93% of the reported PF fell within the measured RF (**Figure 5C**). In both participants, the RF included more hand segments (digits or palmar segments) than did the PF. We verified that this phenomenon was not an artifact of our approach by examining the RFs and PFs characterized independently by another group in a third partially sensate participant using a different characterization method (**Figure 5D**, see Methods and (Herring et al., 2023)). We conclude that the PF of an electrode is determined by the RFs of the neurons around the electrode tip.

**Figure 5.**
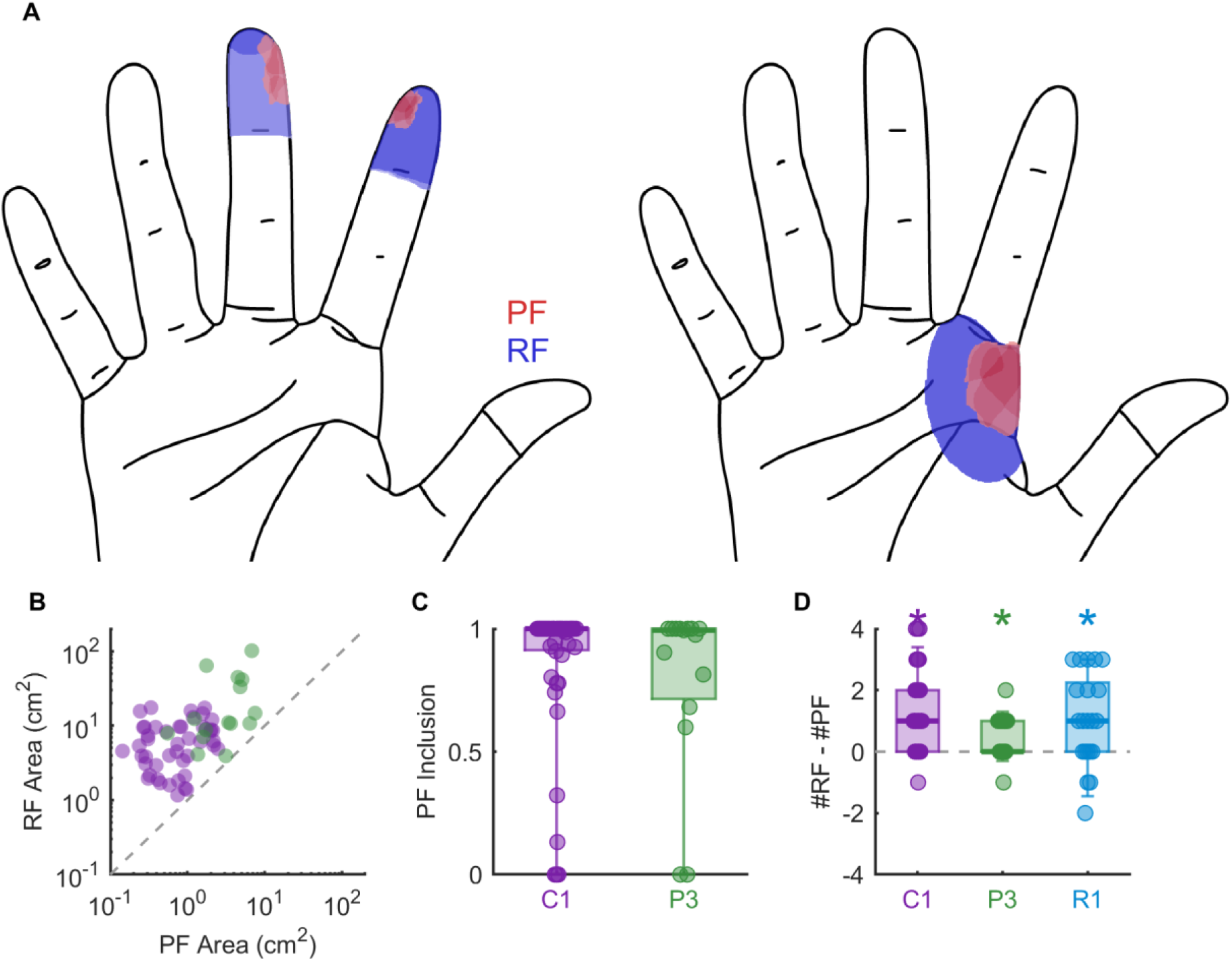
The projected field of an electrode is smaller than and circumscribed by its receptive field. **A**| Aggregate PF (red) and RF (blue) for two example electrodes from participant C1 (left) and P3 (right), respectively. The hue of each denotes the proportion of times a pixel was included in the PF or RF. **B|** Size of the RF vs. size of the PF for electrodes from which both were obtained in participants C1 and P3. Receptive fields were larger for both participants (paired t-test, p < 0.01 for both). Dashed line shows unity. **C|** Proportion of the PF that fell within the RF for all tested electrodes. The median proportion was 1 for both with 25^th^ percentiles of 0.83 and 0.72 for C1 and P3, respectively, suggesting that PFs tended to be completely subsumed by the RF. **D|** Number of regions (digits and palm) encapsulated by each electrode’s RF minus the number of regions encapsulated by its PF for 3 participants (N = 62, 25, 21 for participants C1, P3, and R1, respectively). Multiple electrodes are shown for each participant. RFs spanned more hand regions than PFs in all 3 participants (Wilcoxon rank sum test, p < 0.01, Holm-Bonferroni corrected).

### Projected fields grow larger with more intense ICMS

Next, we examined the degree to which PFs were dependent on ICMS amplitude and frequency. First, we found that the size of the PF increased threefold as amplitude increased from 40 to 80 μA, with ICMS frequency fixed at 100 Hz (**Figure 6A**, median (Q_1_-Q_3_)= 170% (142-330%), 2-way ANOVA, F(2,11) = 6.26, p < 0.01). Second, PFs grew systematically as frequency increased from 50 to 200 Hz, with amplitude fixed at 60 μA (**Figure 6B**, median (Q_1_-Q_3_)= 129% (121-185%), 2-way ANOVA (2,11), F = 7.43, p < 0.01). Because the perceived magnitude increases with both ICMS frequency and amplitude for the electrodes tested (3-way ANOVA (2,2,11), F = 19.61 and 42.35 respectively, p < 0.01 for both), we assessed the relationship between PF size and sensation magnitude. We found that the size of the reported PF could be accurately predicted from the reported magnitude of the evoked sensation, across ICMS frequencies and amplitudes (**Figure 6C**, r = 0.77, p < 0.01). The effect of ICMS intensity on PF size is qualitatively consistent with the proposed neural determinants of the PF. Each volume of cortex maps onto a patch of skin; the greater the volume of cortex activated (amplitude modulation (Kumaravelu et al., 2022; Stoney et al., 1968)) or the number of activated neurons within that cortical volume (frequency modulation (Sombeck et al., 2022)), the greater the corresponding swath of skin.

**Figure 6.**
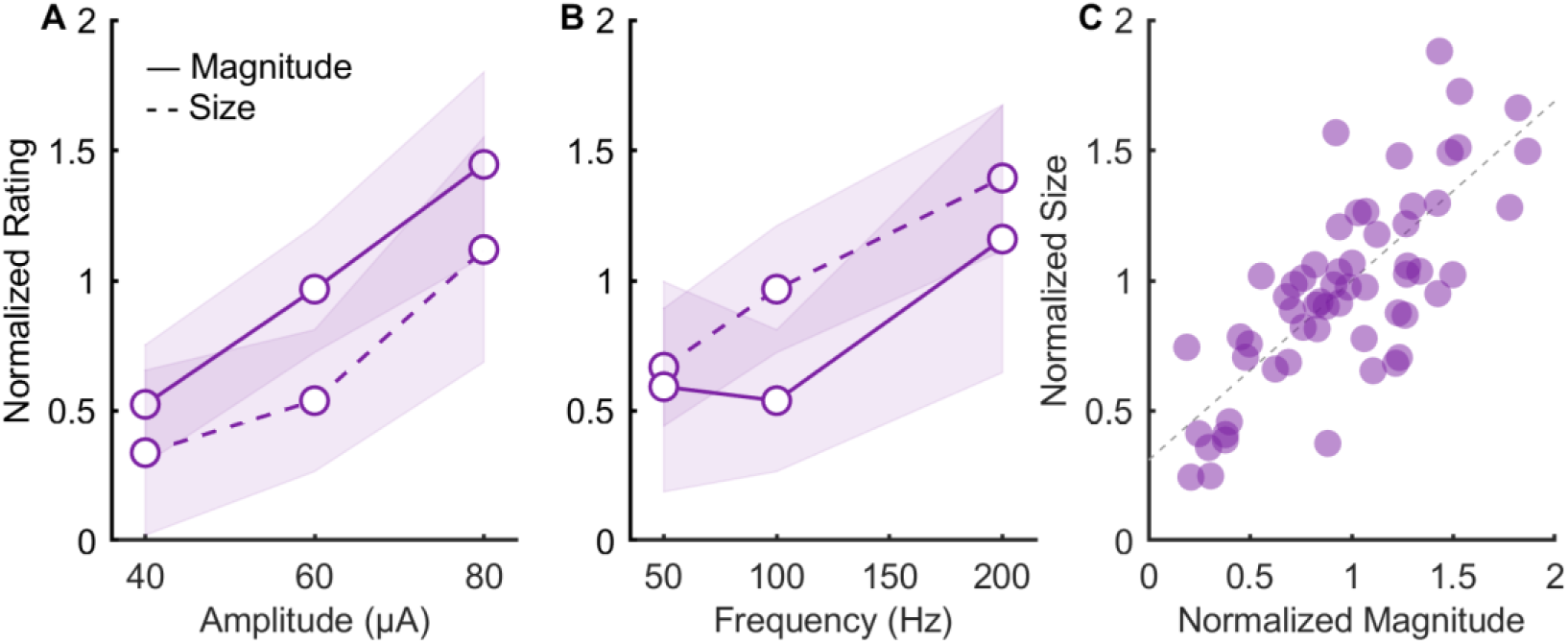
PF size and sensory magnitude increase with ICMS amplitude and frequency. **A**| Normalized ratings of sensory magnitude and PF size vs. ICMS amplitude (with frequency fixed at 100 Hz). The size of the PF for each electrode and condition was normalized by the mean PF across conditions. **B|** Normalized ratings of sensory magnitude and PF size vs. ICMS frequency (with amplitude fixed at 60 µA). Both frequency and amplitude significantly impact perceived size and intensity (2-way ANOVA, p< 0.01 for all). **C|** The effect of ICMS amplitude and frequency on PF size can be accounted for by the latter’s impact on sensory magnitude (r = 0.77, p < 0.01). Data from participant C1.

### Projected fields are superimposed with multi-electrode ICMS

Next, we examined the impact of simultaneously stimulating through multiple electrodes on reported PFs. We wished to assess the degree to which the composite PFs were a conjunction of the component PFs. To this end, we simultaneously delivered ICMS through pairs of electrodes and compared the reported PFs to those when ICMS was delivered through the individual electrodes that formed the pairs (**Figure 7A, B**). In some cases, the electrodes in the pair had PFs that largely overlapped, so we could assess how the two sensations were integrated. In other cases, the electrodes in the pair had non-overlapping PFs, so we could investigate whether the two sensations would interfere or interact with one another. Single and multi-electrode trials were interleaved to limit biases in the PF reports. Examination of the single-electrode and multi-electrode PFs suggests that the latter reflect a superimposition of the former. We tested this by computing the pixel-wise correlation between the sum of the individual PFs and the multi-electrode PF. We found the summed PFs of the component electrodes were highly predictive of the multi-electrode PF (mean r = 0.61, **Figure 7C**). We conclude that PFs combine approximately additively when ICMS is delivered through multiple electrodes.

**Figure 7.**
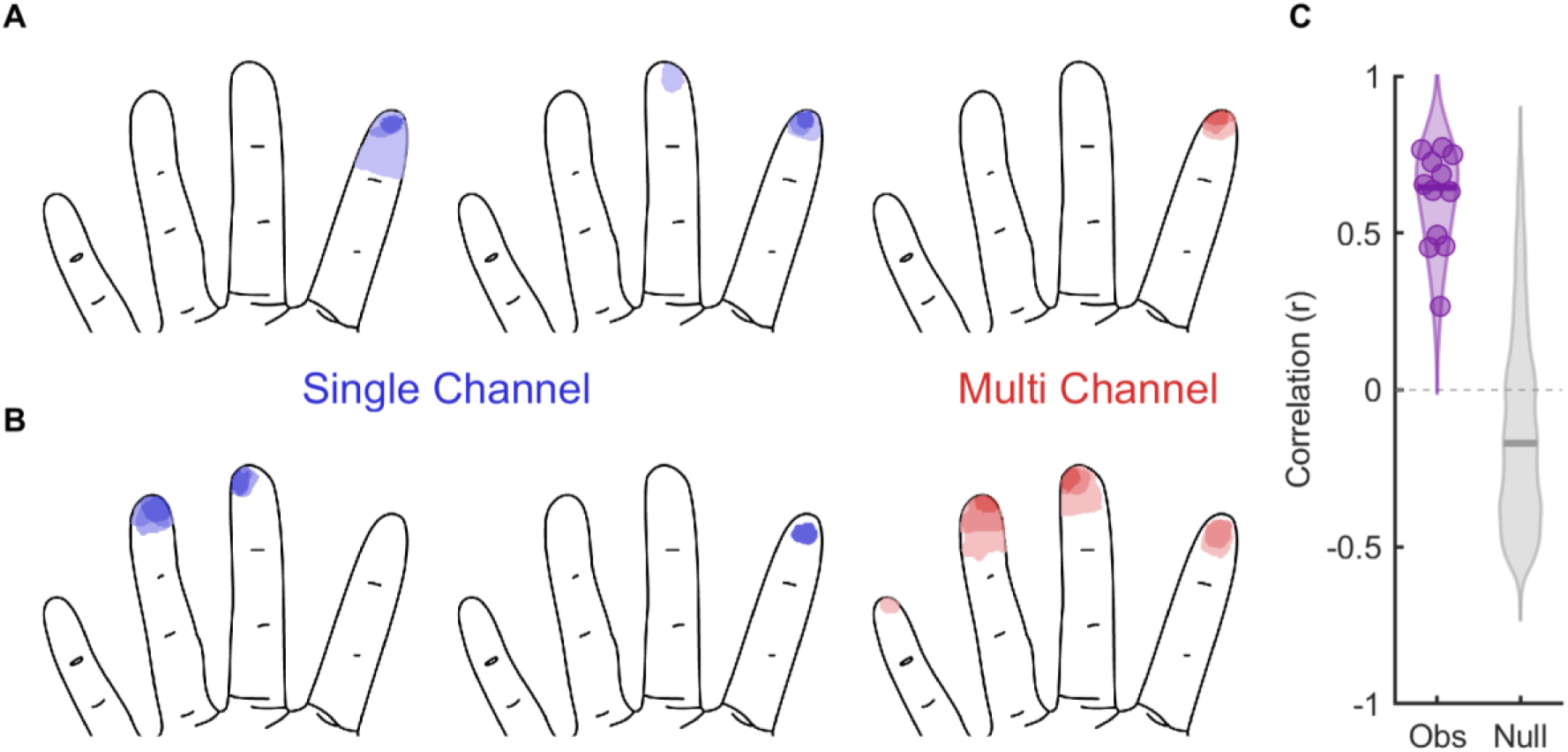
Projected fields with multi-channel stimulation are additive. **A, B**| Two example projected fields predicted from the union of two individual channels versus the reported field when the two channels were stimulated simultaneously. **C|** Correlation between the additive model and observed projected field versus a null model where two random channels were chosen. The additive model significantly outperformed the null model (2-sample Kolmogorov-Smirnov test, D = 0.86, p < 0.01). Data from participant C1.

### Multi-electrode ICMS evokes more localizable sensations than does single-electrode ICMS

As discussed above, the circumscription and systematic localization of PFs can be exploited to convey information about (bionic) hand locations at which contact with an object is established (Flesher et al., 2021, 2016). Given that multi-electrode PFs reflect a superimposition of their component PFs, we reasoned that more focal and thus more easily localizable sensations might be evoked via multi-electrode stimulation. To test this possibility, we mapped force sensors on the bionic hand (Ability Hand, Psyonic) to electrodes with matching PFs. For example, sensors on the prosthetic thumb tip drove stimulation through electrodes with PFs on the thumb tip. In some cases, each sensor drove stimulation through a single electrode; in other cases, sensor output drove a quartet of electrodes (at 60 μA) whose PFs largely overlapped. For these, the overlapping component matched the sensor location. We then randomly touched different bionic digits and had the (blindfolded) participant report which finger was touched (**Figure 8A**). Touches to single digits were randomly interleaved with touches to two digits. We also repeated the same experiment, except that digit(s) were selected by the experimenter on the experimental computer (rather than by physically touching the bionic hand). Results from these two paradigms – one with and one without the bionic hand – were similar and thus combined (**Supplementary Figure 7A**). With either single or multi electrode ICMS, the participant reported the expected digit(s) – corresponding to the PF of the electrode(s) – on the majority of trials. Compared to multi-channel ICMS, however, single-electrode stimulation more often failed to elicit a localizable sensation (30% vs. 0%) and more often elicited a sensation reported on the wrong digit (42% vs. 7%) (**Figure 8A**). The difference in performance for single-vs. multi-electrode stimulation was more pronounced on blocks in which multiple digits were touched, but in both cases multi-electrode stimulation significantly outperformed single-electrode stimulation (Wilcoxon rank sum test, Z = 8.80, p < 0.01 and Z = 9.02, p < 0.01). Multi-channel stimulation thus gives rise to more reliably localizable sensations than does its single-electrode counterpart.

**Figure 8.**
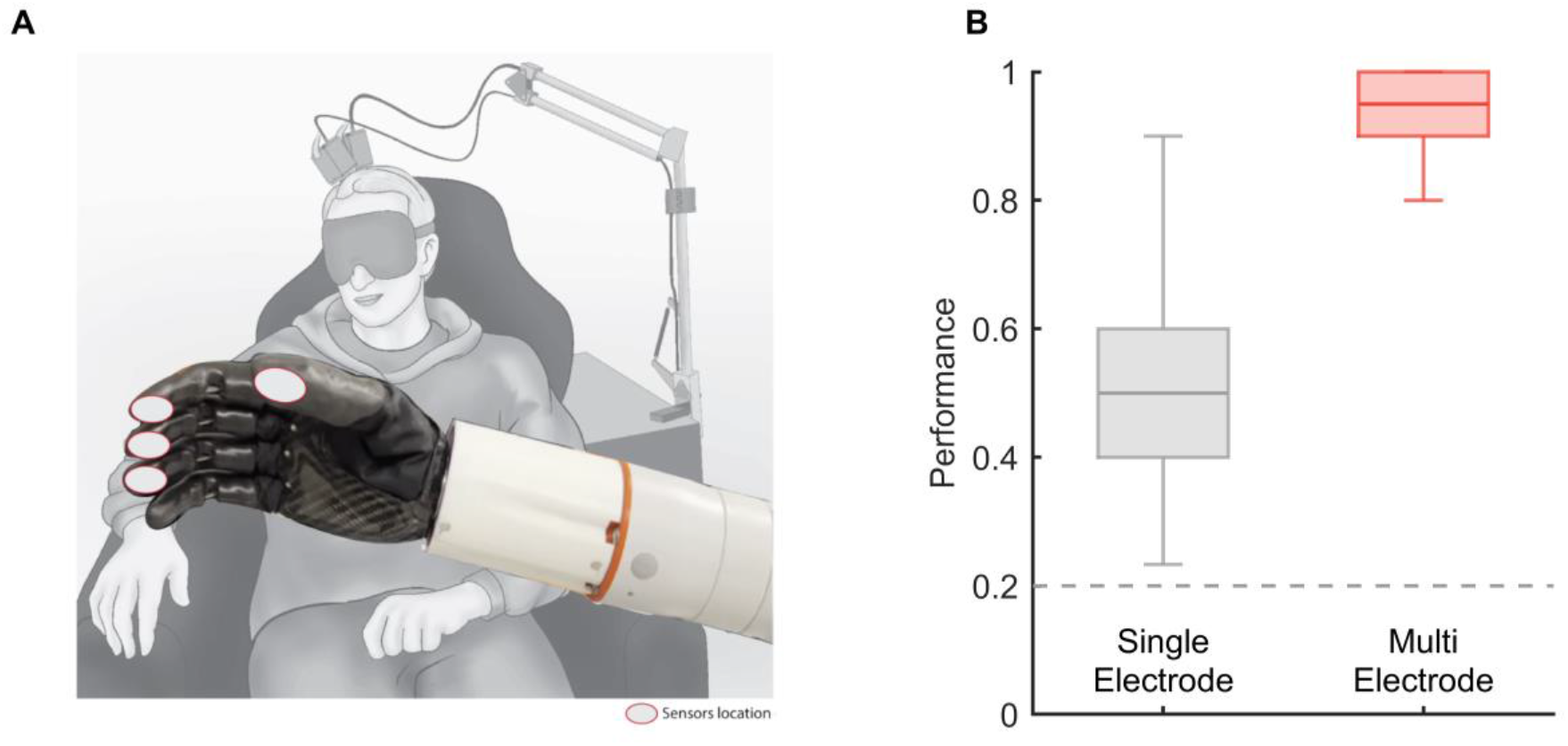
Multi-channel ICMS evokes more localizable sensations than does single-electrode ICMS. **A**| Task setup for robotic digit localization task. The participant was blindfolded while an experimenter randomly squeezed individual prosthetic digits or pairs of digits. **B|** Consolidated performance of robotic and open loop localization tasks. Multi-electrode stimulation evokes more localizable sensations (2-sample t-test, t(236) = 21.6, p < 0.01) Note that trials where the participant failed to detect stimulation are excluded here to reduce confounding localization performance with detectability. Data from participant C1.

## Discussion

### PFs are highly stable over time

PFs remained stable over the testing period, which spanned ∼2 years for participants C1 and P3 and ∼7 years for participant P2. Indeed, the variability in the PF overlap and centroid distance across repeated tests on the same electrode were equivalent within and across sessions. Given that the PF location largely coincides with RF location, this stability suggests that the somatosensory homunculus is highly stable, consistent with fMRI studies with amputees (Kikkert et al., 2016) and people with tetraplegia (Andersen and Aflalo, 2022; Kikkert et al., 2021). Moreover, the participants varied in their residual sensations, where participants C1 had spared sensation over most of his hand, P3 over all but the little finger, and P2 only on the thumb and index. Across participants, the PFs were indistinguishable in their size and relative locations whether these were on sensate or insensate patches of skin, suggesting that the homunculus remains stable regardless of the degree of residual sensation. Furthermore, recent results with intraneural electrodes implanted in the human nerves suggested that the PF changes observed over time are most likely caused by the foreign body reactions and their impact on the coupling between nerve and electrode (Valle et al., 2022). Indeed, deliberate attempts to modify the PFs of electrodes implanted in the nerves – which leverage any subcortical plasticity – failed to systematically change the PF location (Ortiz-Catalan et al., 2020a), further evidence for the stability of body maps in adulthood (Makin and Bensmaia, 2017).

### The projected field of an electrode is subsumed by its receptive field

Since the PF of an electrode is determined by its location on the somatosensory homunculus, the spatial extent of the PF is expected to be determined by the spatial extent of activation across the cortical surface. Consistent with this, increases in ICMS amplitude or frequency lead to increases in the volume of activated neurons and/or density of activated neurons within a volume (Dadarlat et al., 2022; Kumaravelu et al., 2022; Sombeck et al., 2022; Stoney et al., 1968) and concomitant increases in PF area. Furthermore, the PF of an electrode is, on average, a sixth to a seventh the size of its RF. If we assume that the spatial extent of a PF is determined by the activated cortical area, ICMS at 60 μA activates the somata of neurons over a span of around 1 mm (Kumaravelu et al., 2022) (**Supplementary Figure 8A**). The RF size is determined by (1) the area over which neurons contribute to the multiunit activity on a given electrode (∼150 μm radius) and (2) the area of cortex that is activated by a touch (**Supplementary Figure 8B**). Regarding the latter, a punctate indentation applied to the skin evokes neuronal activity on two spatial scales (**Supplementary Figure 8C**). Contact initiation evokes a transient response over a wide swath of cortex (spanning ∼15 electrodes on the array), with RFs spanning multiple digits. Contact maintenance is associated with a much more spatially restricted sustained response (spanning ∼2 electrodes). RFs, mapped by touching the skin lightly and repetitively, reflect the transient response. If responses to contact transients determined the spatial extent of a tactile experience, a touch on a single digit would evoke a sensation on multiple digits. Accordingly, the spatial extent of a tactile sensation is more likely determined by the cortical population that remains active during maintained contact, the so-called hotzone (Callier et al., 2019). For a punctate indentation, the hotzone spans 1 to 2 electrodes. At a first approximation, then, the area of cortex activated by ICMS is on the same order of magnitude as the hotzone, and much smaller than the transiently activated cortical area (**Supplementary Figure 8D**). Note, however, that the spatial extent of a sensation is not only determined by the spatial extent of the activated cortex but also by the spatial pattern of that activation. A punctate indentation produces a response that is spatially graded, peaking at the hotzone and dropping off smoothly around it (Callier et al., 2019). While the ICMS-evoked activation also decreases with distance from the electrode tip (Merrill et al., 2005), the pattern is almost certainly different. This difference likely underlies the fact that ICMS-evoked sensations have more diffuse borders than do their natural tactile counterparts. The relationship between the spatial pattern of cortical activation and the evoked sensation remains to be conclusively elucidated.

### Multi-electrode stimulation evokes more localizable sensations

Mapping sensors on the bionic hand to somatotopically matched electrodes in S1 yields intuitive feedback about the locations on the bionic hands at which contacts with an object occur. Indeed, contact with the prosthetic thumb tip, for example, leads to a sensation experienced on the thumb tip. Here, we show that this mapping is even more effective if the sensor output drives ICMS through multiple electrodes, whose common PF matches the sensor location. For example, if multiple electrodes have PFs that include the thumb, their aggregate PF will be dominated by the thumb. Multi-electrode ICMS thus results in sensations that not only are more tangible but also more readily localizable to the hand regions corresponding to the PF overlap. While the same outcome might be achieved by increasing the amplitude of ICMS delivered through a single electrode, resulting in a more salient PF core despite the concomitant increase in PF area, multi-channel ICMS enables improved localizability within the safety limits of ICMS, capped at 100 μA (Rajan et al., 2015). In addition to the improvement in localization, multi-channel ICMS also leads to an improvement in force feedback compared to its single-channel counterpart, as evidenced by a greater number of discriminable steps of force within the safe and perceptible range of amplitudes (Greenspon et al., 2023). Moreover, multi-channel ICMS robustly evokes somatosensory sensations at significantly shorter latencies than does single-electrode ICMS (Bjånes et al., 2022; Sombeck and Miller, 2019). On the other hand, multi-electrode ICMS of S1 has a stronger deleterious effect on the ability to decode motor intent by evoking more activity in M1 than does single-electrode ICMS (Shelchkova et al., 2022). Indeed, the intermixing of motor and sensory signals is liable to disrupt the decoders, particularly if these are not trained in the presence of ICMS-based feedback. This disruption, however, can be minimized by implementing biomimetic feedback – which emphasizes contact transients at the expense of maintained contact (Bensmaia et al., 2020; Okorokova et al., 2018). Overall, then, multi-electrode stimulation leads to more effective tactile feedback for bionic hands.

### Implanted arrays should consist of distributed clusters of electrodes

As the majority of our interactions with objects involve the digit tips (Christel, M.I. et al., 2020), ICMS-tactile feedback emphasizes fingertip sensations over more proximal ones. Implants in S1 often consist of 6×10 arrays of electrodes with every other electrode wired in a checkerboard pattern (**Figure 1**). The placement of the arrays is based on the identification of distinct digit-specific patches of activity using fMRI or MEG. Individual arrays are then placed near the border of two digits to maximize coverage.

While the aforementioned imaging technologies yield reliable localization of the individual digit representations in S1 (e.g. digit 2 vs. digit 3), localization of the digit tips (vs the digit bases) is far less clear. In fact, the location of the digit tip representation in postcentral gyrus is unclear. In humans and monkeys, RFs progress from digit tip to base as one progresses caudally through Brodmann’s area 3b from its border with area 3a (Blankenburg et al., 2003; Delhaye et al., 2018; Rosa M. Sanchez-Panchuelo et al., 2012; Roux et al., 2018). The RF progression reverses at the border with area 1, proceeding from base to tip, then reverses again at the border with area 2. Given that digit tip representations in area 3b are deep in the central sulcus, the only access to digit tip representations is at the area 1/2 border with the arrays currently approved for human use, which cannot access deep structures. As much of area 2 may be in the anterior bank of the intraparietal sulcus (IPS) in humans (Geyer et al., 1999), the digit tip representations are likely to be near the IPS, though this does not seem to always be the case. While implanting larger arrays may improve the coverage, implanting more arrays rather than larger ones might improve the chances that all digits are represented. Ideally, clusters of electrodes (4 or 9 electrodes) would be implanted in the center of the presumptive digit tip representations, near the IPS (**Supplementary Figure 9**). With an inter-electrode spacing of 400 μm, the clusters would activate populations of neurons with largely overlapping RFs yet different electrodes would activate largely non-overlapping neuronal populations as the electric fields at each electrode would be largely nonoverlapping over the range of safe ICMS amplitudes. This configuration can scale up as the robotics, sensorization, and algorithms improve, with each cluster matched to a sensor on the hand.

## Methods

### Participants

This study was conducted under an Investigational Device Exemption from the U.S. Food and Drug Administration and approved by the Institutional Review Boards at the Universities of Pittsburgh and Chicago. The clinical trial is registered at ClinicalTrials.gov (NCT01894802). Informed consent was obtained before any study procedures were conducted. Participant C1 (m), 57 years old at the time of implant, presented with a C4-level ASIA D spinal cord injury (SCI) that occurred 35 years prior to implant. Participant P2 (m), 28 years old at the time of implant, presented with a C5 motor/C6 sensory ASIA B SCI that occurred 10 years prior to implant. Participant P3 (m), 28 years old at the time of implant, presented with a C6 ASIA B SCI that occurred 12 years prior to implant. Participant R1 (m), who performed a subset of experiments, was enrolled under a separate IRB and IDE-approved clinical trial (NCT3898804). R1 presented with a C3/C4 ASIA B SCI that occurred 5 years prior to implant, and his cortical implants are described in detail in (Herring et al., 2023).

### Residual sensation

Participant C1 had retained sensation over the entire volar surface of the hand, with nearly normal detection thresholds (measured with van Frey hairs) on the digit tips, and elevated thresholds over the rest of the hand (**Supplementary Figure 6**). Participant P2 had retained sensation over the thumb, index finger, and underlying palm, albeit with elevated thresholds, but the rest of his hand was insensate. Participant P3 had nearly normal thresholds on the thumb and most of the index finger, higher thresholds on the middle finger, was nearly insensate on the ring finger and completely insensate on the little finger.

### Cortical implants

We implanted four microelectrode arrays (Blackrock Neurotech, Salt Lake City, UT, USA) in each participant (C1, P2, P3). The two arrays (one medial and one lateral array) in Brodmann’s area 1 of S1 were 2.4 mm × 4 mm with sixty 1.5-mm long electrode shanks wired in a checkerboard pattern such that 32 electrodes could be stimulated. The two arrays in primary motor cortex were 4 mm × 4 mm with one-hundred 1.5-mm long electrode shanks wired such that 96 (C1 and P3) or 88 (P2) electrodes could be used to monitor neural activity. The inactive shanks were located at the corners of these arrays. Two percutaneous connectors, each connected to one sensory array and one motor array, were fixed to the participant’s skull. We targeted array placement during surgery based on functional neuroimaging (fMRI) or magnetoencephalography (MEG) of the participants attempting to make movements of the hand and arm (all participants) and imagining feeling sensations on their fingertips (participant P2), within the constraints of anatomical features such as blood vessels and cortical topography. Participant R1 was implanted with two 64-channel arrays (8 × 8) in Brodmann’s area 1, with PFs primarily on the index and middle fingers (see (Herring et al., 2023) for details).

### Intracortical microstimulation

ICMS was delivered via a CereStim 96 (Blackrock Neurotech). Each stimulating pulse consisted of a 200-µs cathodic phase followed by a half-amplitude 400-µs anodic phase (to maintain charge balance), the two phases separated by 100 µs. For multi-channel stimulation, the same pulse train was delivered to all channels simultaneously and synchronously.

### Projected field mapping and quantification

To map a PF, ICMS was delivered through one or more electrodes (60 µA, 100 Hz, 1 second unless otherwise specified), and the participant reported whether a sensation was evoked. The participant could request as many repetitions of the stimulus as desired. During or after the stimulation, the participants drew the spatial extent of the sensation on a digital representation of the hand using a stylus.

Periodically (every 1-3 months), this procedure was repeated for every electrode on both sensory arrays, with electrodes interleaved in random order, across a span of approximately 800, 2750, and 750 days for participants C1, P2, and P3, respectively. From these digital images, we counted the number of pixels subsumed by the PF, computed its area and centroid (center of mass). To gauge the validity of the computed center of mass, one participant also marked, on a subset of electrodes in one session, the center of the PF on the picture of their hand. We found that the estimated centroids matched the reported centroids with sub-millimeter accuracy (median = 0.71 mm, Q_1_, Q_3_ = 0.52, 1.33 mm).

In a fourth participant (R1), the PF mapping procedure differed slightly. On each experimental block, a group of 5 electrodes were randomly selected, and each electrode was stimulated approximately 5 times (n = 4-7 times, mean = 5.12) in randomized order within the block. The participant verbally reported the hand segments on which the sensation was experienced (distal and medial finger pads, e.g.), each segment numbered on a digital representation of the hand (Herring et al., 2023). ICMS was identical as that for the other participants, except that pulse amplitude was 80 µA. Skin locations included in the reported PF on at least 60% of trials for a given electrode were included in its PF.

### Aggregate projected fields

Having established that day-to-day variations in PF location were essentially random and primarily reflected noise in the reports, we constructed an estimate of each PF that weighted skin regions according to the frequency with which these were included in the single-day PF reports. Specifically, we computed the proportion of times a given pixel was included in the single-day PF. We then eliminated pixels that were included on fewer than 25% of single-day reports and normalized the remaining proportions such that they summed to one. For some electrodes, this criterion eliminated all of the pixels. These electrodes were not included in subsequent analysis and are reported as having no reliable PF. In the computation of the aggregate centroid, each pixel was weighted according to its value (normalized proportion). When aggregate centroids spanned multiple hand segments, the one with the largest summed pixel values was identified as the dominant one. The size of the aggregate field was the total number of pixels that exceeded the threshold.

### Projected field stability

We gauged projected field stability in three ways. First, we computed the distance between each single-day PF centroid and the centroid of the first reported PFs. Because this measure was susceptible to an anomalous report on the first session, we also computed the distance between the PF centroid on any given session and the centroid of the aggregate PF. Third, we measured the proportion of all unique pixels reported for a given electrode that exceeded each of several thresholds, reasoning that stable PFs would be consistent across a wide range of thresholds.

### PF progression over the cortical surface

To test the degree to which PF progressions follow the canonical homunculus, we assessed how well we could identify the dominant hand segment for an electrode given that electrode’s location. In brief, we used the row and column coordinates of the array as the X and Y coordinates in Cartesian space, whose origin was the center of the array. We then projected the coordinates onto each of 72 axes, spanning the range from 0 to 360 degrees in increments of 5 degrees. For example, for the 0-degree axis, the coordinate would be identical to the X coordinate (row). For each axis, we then applied a linear discriminant classifier with the projected coordinates as inputs and the dominant hand segments (digit, e.g.) as output classes (MATLAB’s *fitcdiscr*). To further test the degree to which PFs progress somatotopically, we computed, for each pair of electrodes, the distance between their respective aggregate centroids and assessed the relationship between PF centroid distance and the physical distance between the electrodes over the cortical surface.

### Receptive field mapping

To map the extent of the receptive fields of the multi-unit around each electrode, we lightly brushed the skin of the hand using cotton swabs or Von Frey hairs while monitoring the multi-unit response using a speaker. One experimenter stimulated the hand while the other experimenter selected the electrode to be mapped (unknown to the first). In the case of participant C1, where 3 sessions were completed, the electrode order was randomized across sessions. The RF was reported on the same tablet interface used for PFs so that the same analytical protocols could be applied to both types of fields. An aggregate RF was produced for participant C1 with a cutoff of 33% (the pixel must have been present on 1/3 of observations) using the method described above for PFs. Measured RFs were nearly identical across sessions in the one participant for whom these were measured repeatedly over multiple sessions. For participant R1, a trapezoidal indentation was delivered to the distal finger pad of each digit using a tactile linear actuator (LCA25-025-6MSD6, SMAC, Carlsbad, California, USA) while recording the neuronal activity evoked in S1. Each indentation was presented 60 times at a constant amplitude. Because the neuronal signal was weak, we computed spike band power – the root mean square of the filtered signal in 1 ms bins – to quantify the strength of the neural response. The spike band power was normalized within trials by z-scoring the signal during the 2 second stimulus by the 2 second pre-stimulus interval, then the normalized metric was trial averaged for each indenter location. Electrodes with peak normalized spike band power greater than 1.5 standard deviations above baseline were considered responsive to indentation at the specific location. While this RF mapping approach involved a much more controlled and systematic mechanical stimulus than that used in the other three participants (P2, P3, C1), it suffers from the poor coverage, able only to detect neuronal responses evoked by stimulation of the distal digit tips.

### Localization task

To assess the degree to which ICMS conveys information about the location of contact between bionic hand and objects, we implemented a sensory encoding algorithm that linked sensors on the robotic hand (Ability Hand, Psyonic, CA, USA) and somatotopically matched electrodes in S1. For example, the output of a sensor on the index fingertip drove ICMS delivered through one or more electrodes with PFs on the index fingertip. Then, on each trial, the experimenter squeezed one or more digits and the participants task was to report which digits were squeezed. Single-digit and two-digit trials were randomly interleaved and the participant was warned that one or two fingers might be squeezed on any given trial. On some experimental blocks, the digit(s) were selected by the experimenter, and an automated protocol randomly selected stimulating channels corresponding to the selected digits for testing, allowing for greater flexibility to test different channels. Sensor output drove ICMS (at 100 Hz) through a single electrode with a PF on the corresponding digit or through multiple electrodes, all with PFs that were predominantly on that digit. The amplitude of the ICMS was proportional to the sensor output and capped at 60 μA; for multi-electrode stimulation, the same ICMS was delivered to all the channels simultaneously. For computercontrolled trials, selection of digit triggered a 1-sec, 100-Hz, 60-μA ICMS pulse train delivered through 1 or 4 electrodes. We could then compare the participant’s performance with single vs. multi-channel stimulation on the single-vs. multi-digit task. The participant reported the location of the sensation (individual digits, combinations of digits, or no sensation) at the end of the stimulus. On single-digit trials, the participant received full credit if he correctly identified the digit where the aggregate PF was located; on multi-digit trials, he received half credit for each correctly identified digit.

## Acknowledgments

We would like to thank the participants for their generous contribution to the advancement of science. The work at the University of Chicago and University of Pittsburgh was supported by NINDS grants UH3 NS107714, R35 NS122333 and RO1 NS119160. The work at Case Western Reserve University was supported by grants CDMRP SCIRP SCI80308 and VA I01RX002654.

## Disclosures

NH and RG serve as consultants for Blackrock Neurotech, Inc. RG is also on the scientific advisory board of Neurowired LLC. MB, JC, and RG received research funding from Blackrock Neurotech, Inc. though that funding did not support the work presented here.

## Supplementary Figures

**Supplementary Figure 1.**
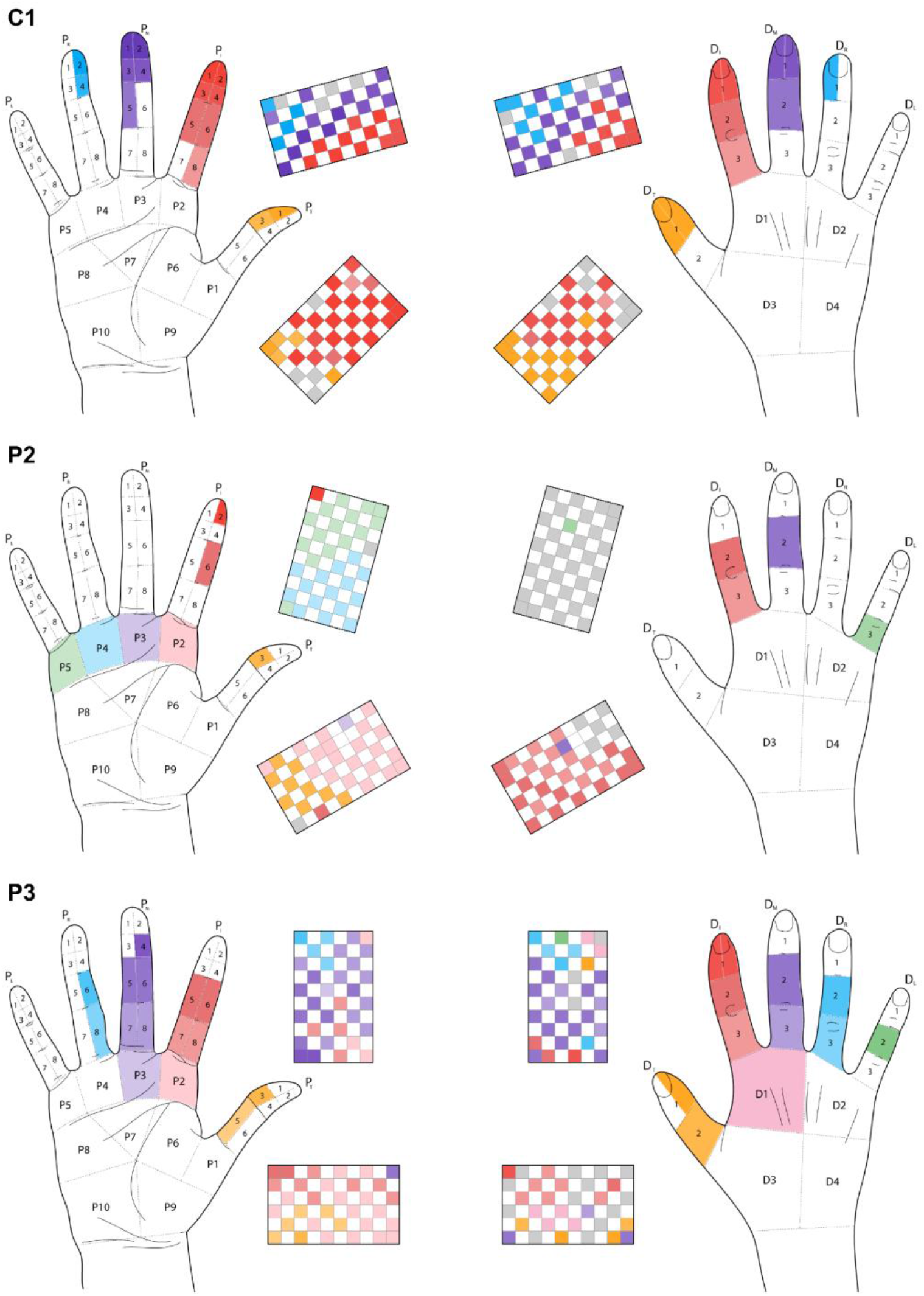
Palmar and dorsal projected field maps for all participants. The hand diagrams show the distributions of the locations of the sensations evoked by ICMS any electrode on the palmar (left) and dorsal (right) surface of the hand. The array diagrams show the dominant hand segment for each electrode. All array rotations are approximately aligned such that up is medial and anterior is left.

**Supplementary Figure 2.**
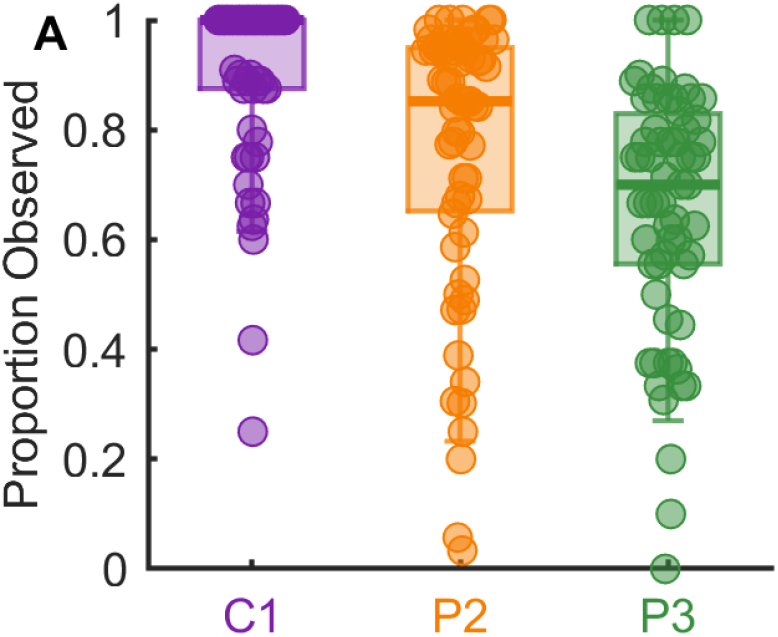
Incidence of evoked sensations. Proportion of times that ICMS (at 100 Hz, 60 µA) through a given electrode evoked a sensation in the three participants.

**Supplementary Figure 3.**
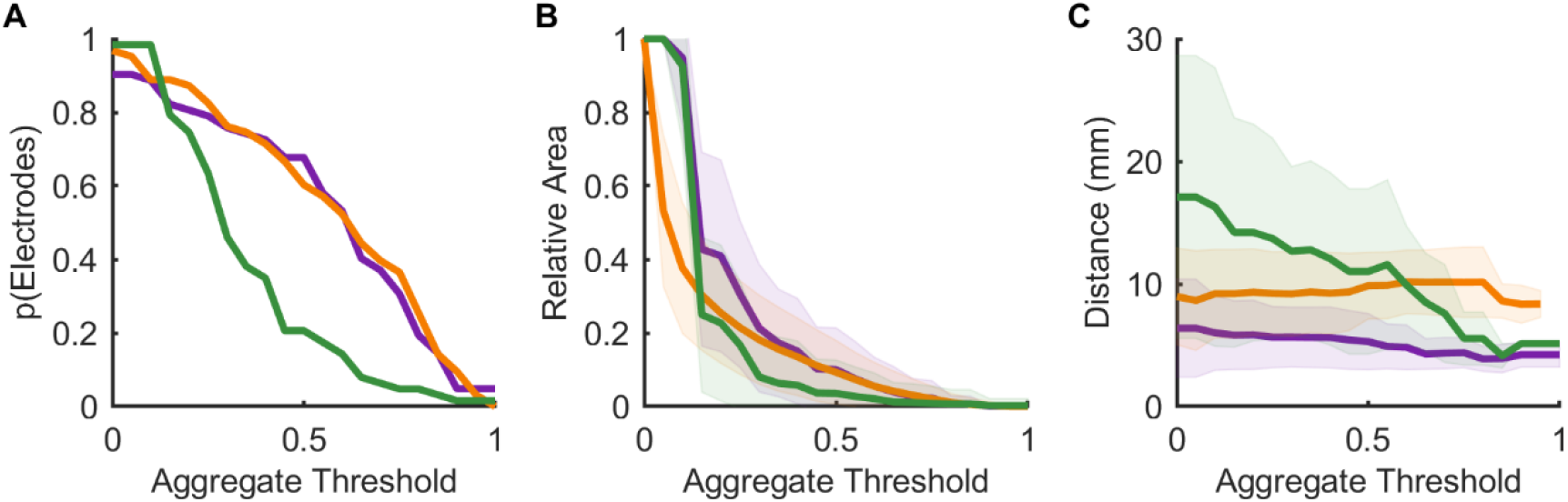
Thresholding the aggregate PFs to assess stability. **A**| The proportion of electrodes for which any pixel exceeds the criterion decreases as the threshold increases. **B|** The area of the aggregate PF drops dramatically as the threshold increases to 0.25 and then decrease more slowly. **C|** The variability in the PF (Euclidean distance of each observation’s centroid from the aggregate centroid) changes only marginally as threshold increases.

**Supplementary Figure 4.**
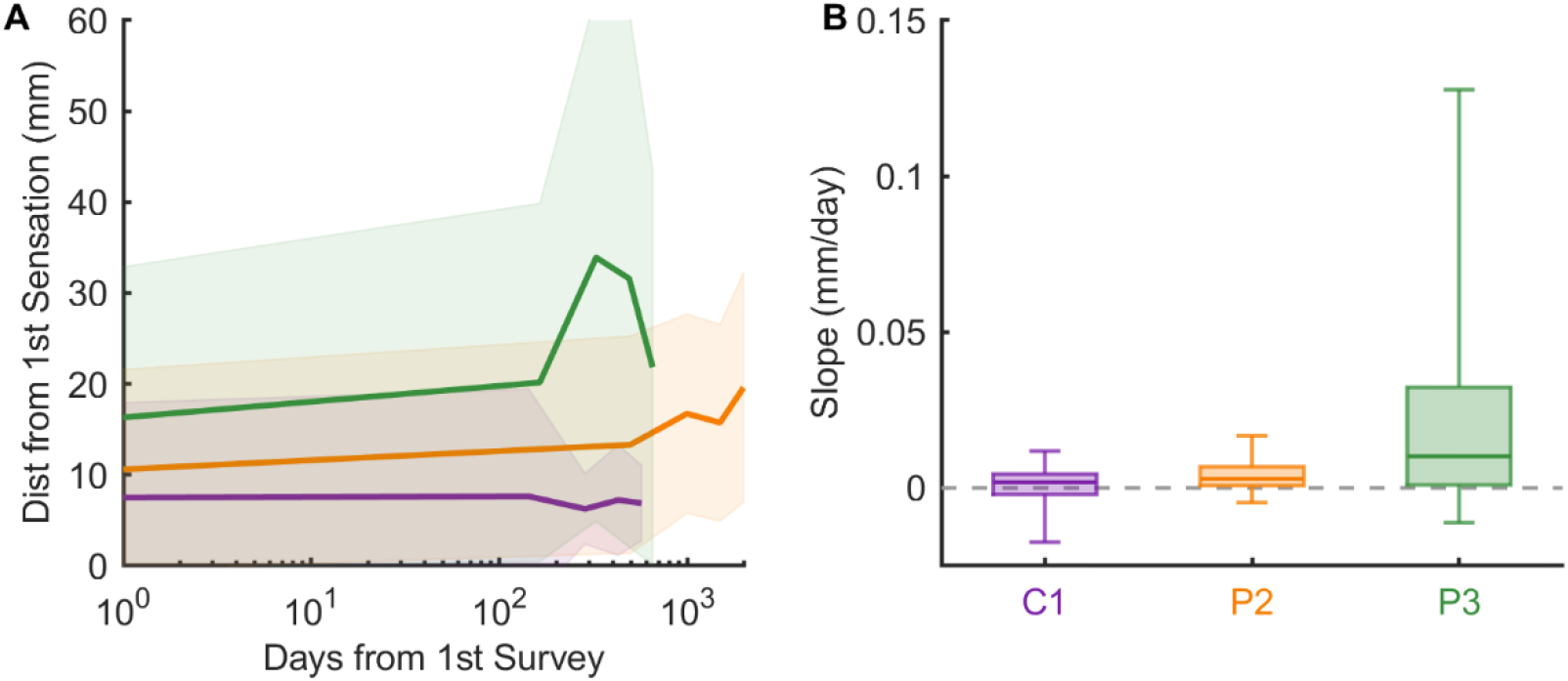
Sensations are stable over time. **A**| The mean distance between the first centroid and subsequent ones is stable. **B|** The distribution of slopes computed from the data shown in panel A for individual electrodes grouped by participant.

**Supplementary Figure 5.**
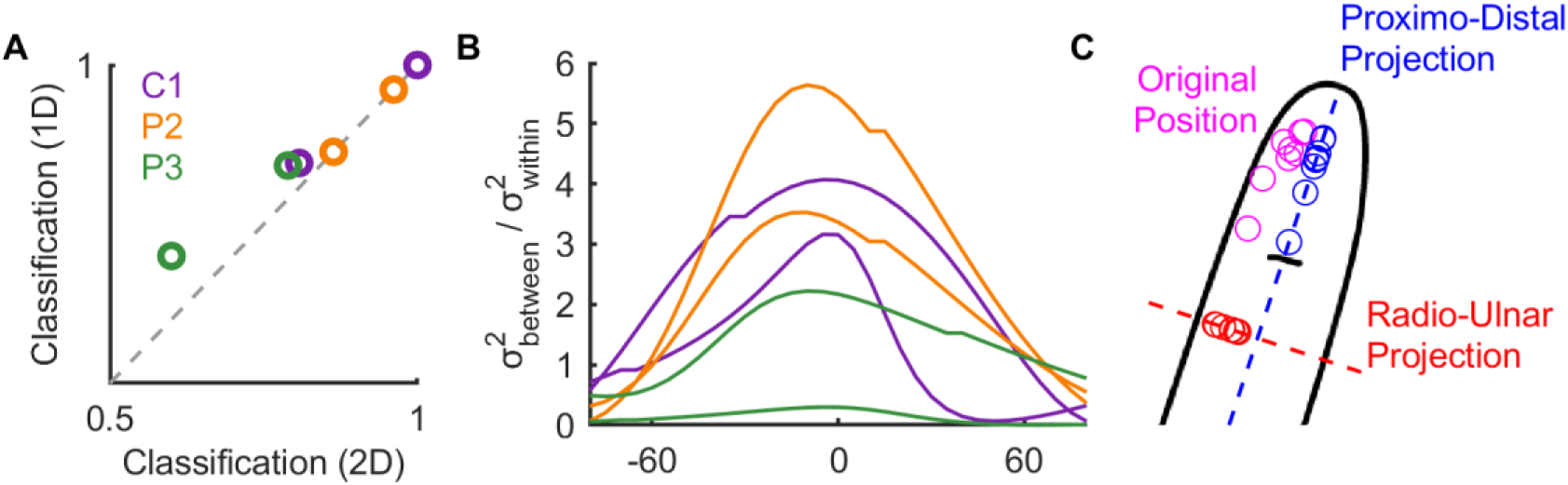
Percepts are distributed along anatomical axes within S1. **A**| Digit classification when using either both axes (row & column) or the best single projected axis. **B|** Within class variance divided by across class variance for digit discrimination LDA as a function of rotation relative to the central sulcus. **C|** Example projections of percept centroids in their original position (magenta) when projected along either the proximo-distal (blue) or radio-ulnar (red) axes.

**Supplementary Figure 6.**
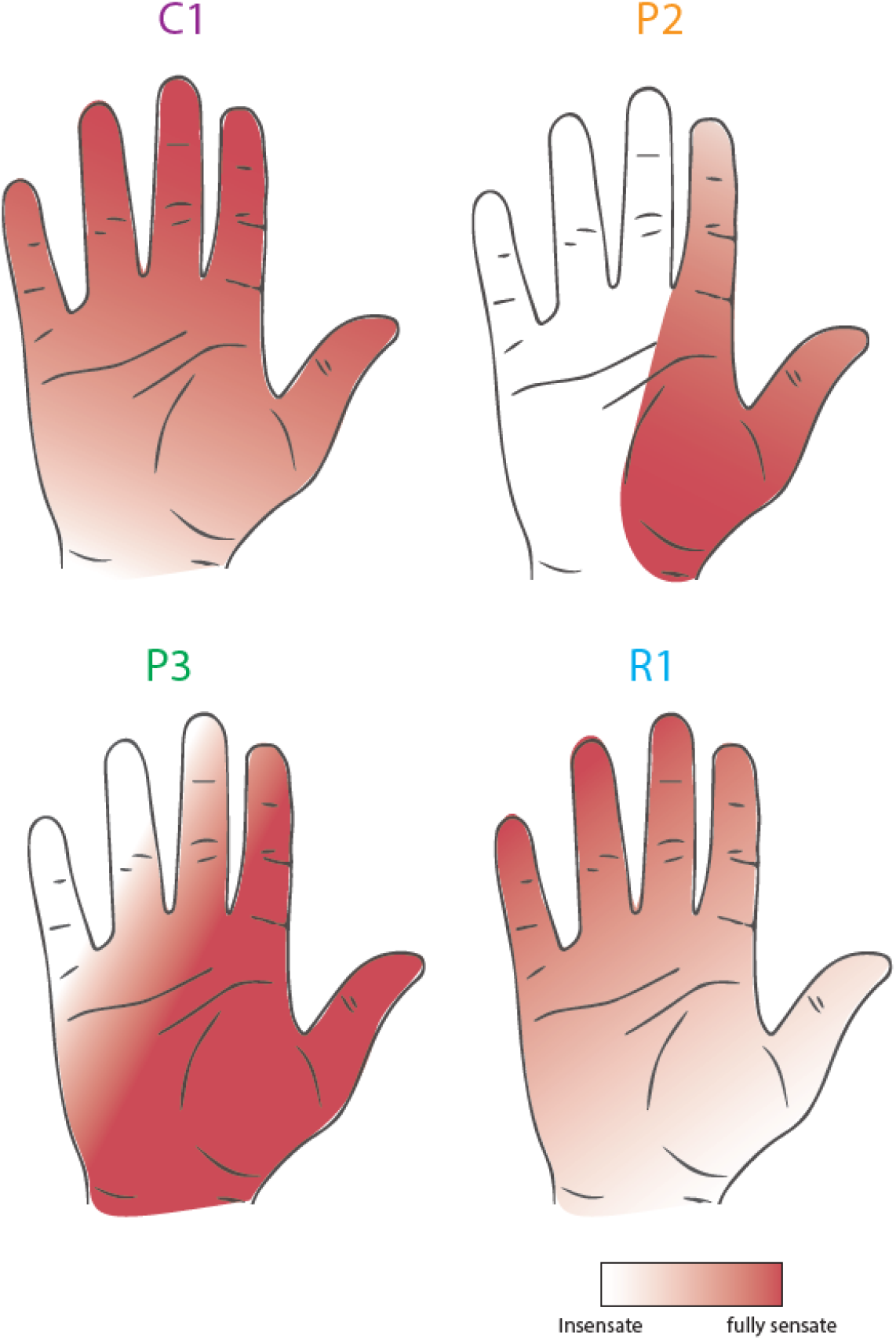
Residual touch sensation for each participant.

**Supplementary Figure 7.**
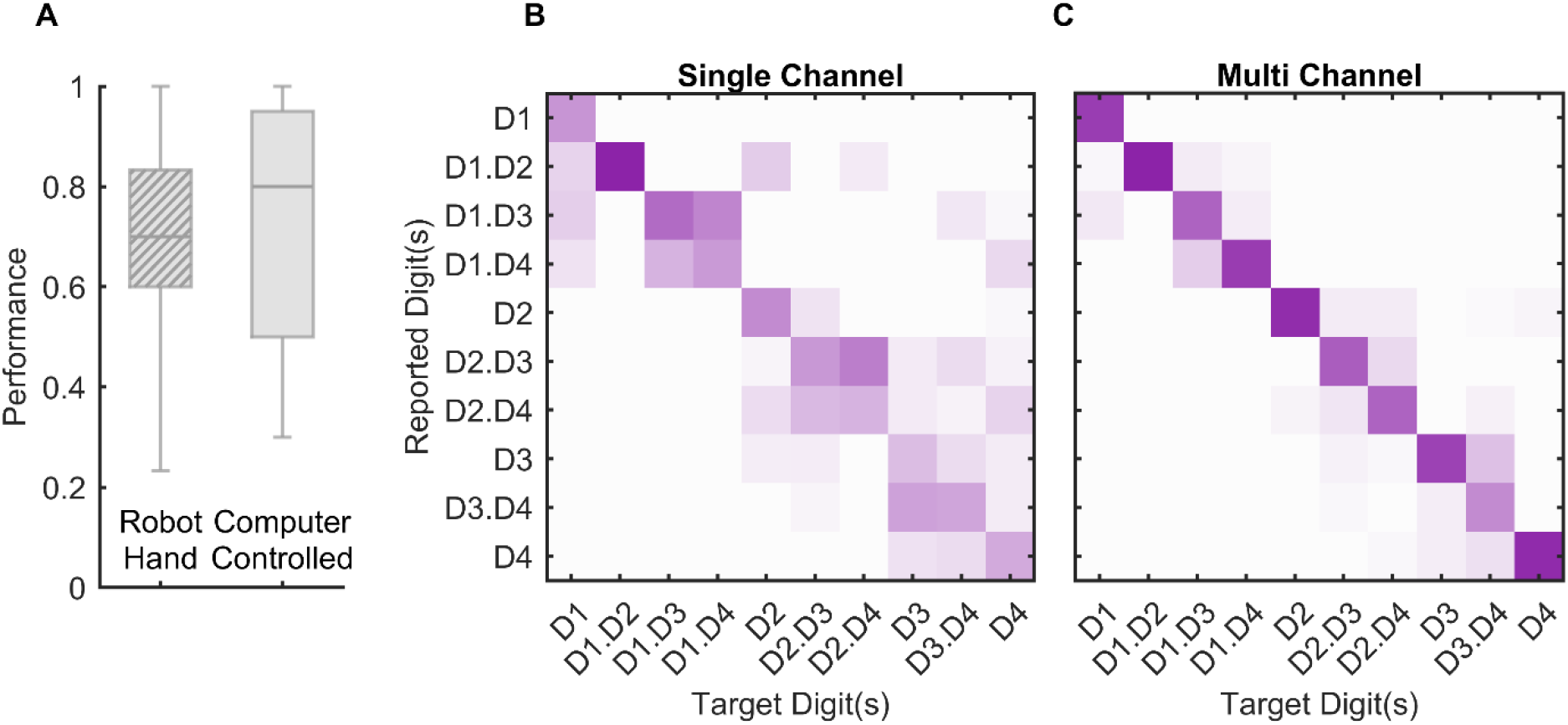
Multi-electrode ICMS leads to improved localization. **A**| Performance on the localization task when stimuli were triggered via the robotic hand or the computer. Overall performance was statistically equivalent (2-sample t-test, t(236) = 0.58, p = 0.56). **B|** Confusion matrices for single and multi-channel stimulation. Data from participant C1.

**Supplementary Figure 8.**
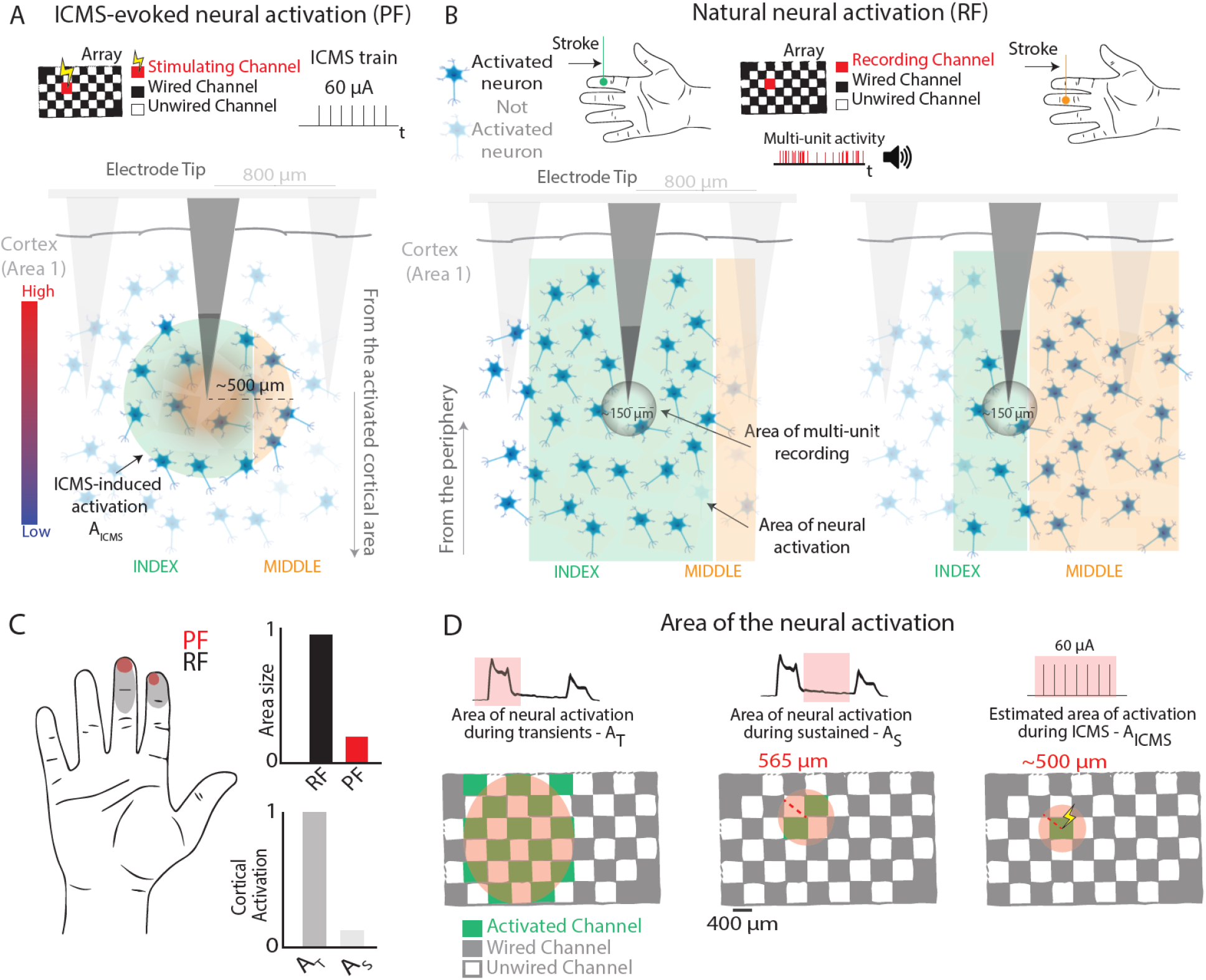
Cortical activation during natural touch and ICMS. **A**| Illustration of cortical activation by ICMS and **B**| during RF mapping for a given electrode. The cortical activation by touch is much wider than that by ICMS. **C**| Considering the differences in cortical activation during transient and sustained phases of contact, the area of cortex activated by ICMS (A_ICMS_) is on the same spatial scale as the hotzone (cortical activation during sustained contact - A_S_), but systematically smaller than the transiently activated cortical area (-A_T_), at a first approximation.

**Supplementary Figure 9.**
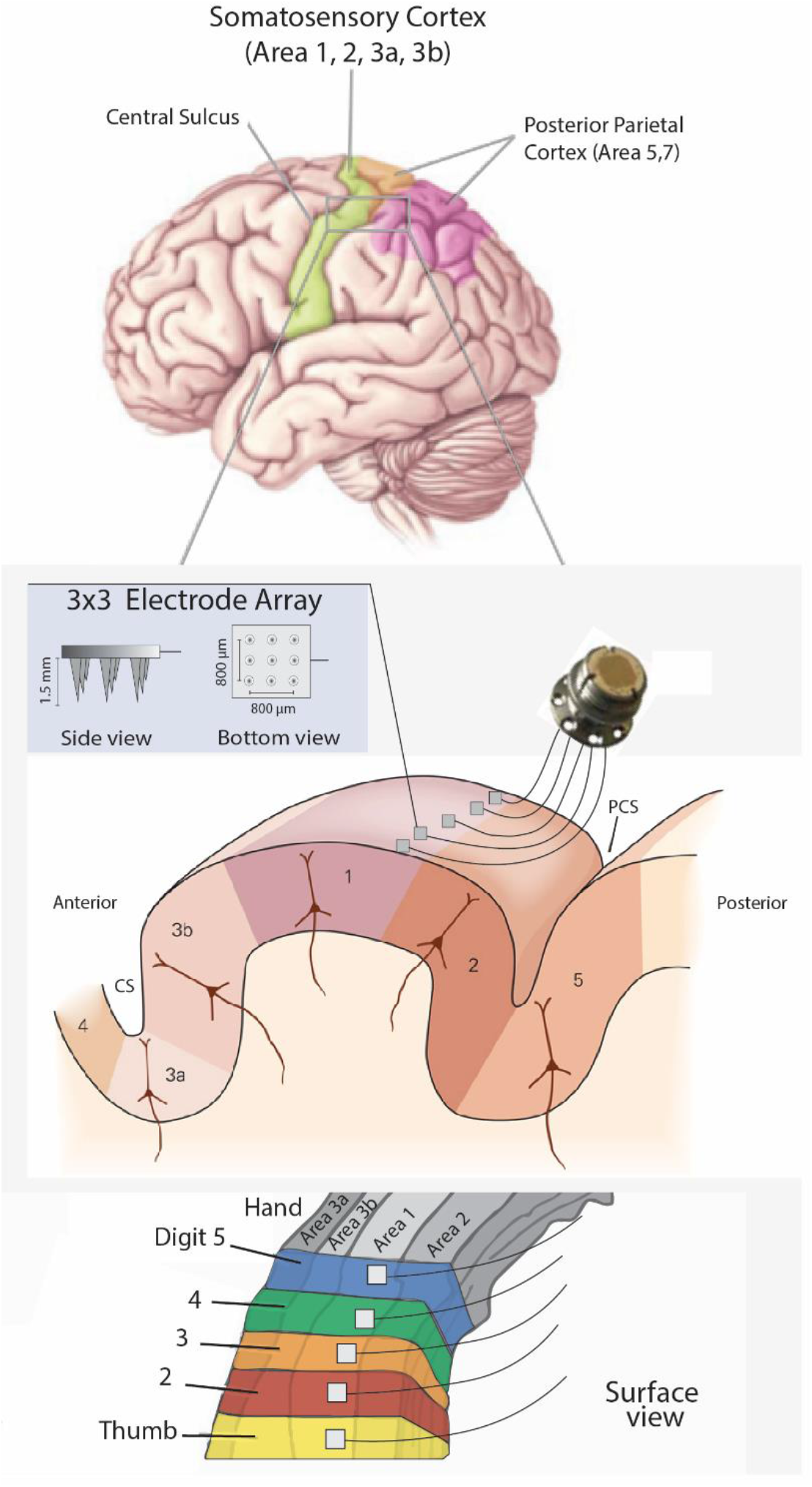
Proposed design of the neural interface. Five 3 × 3 arrays are implanted in Brodmann’s area 1 near its border with area 2, each centered on the respective representations of the five digits, identified using fMRI, MEG, or ECog.

## Notes

### Summary of Updates

Fixed some errors in the calculation of the PF and RF area and made minor edits throughout.

